# Signalome-wide assessment of erythrocyte response to *Plasmodium* reveals novel targets for host-directed antimalarial intervention

**DOI:** 10.1101/2020.06.09.143206

**Authors:** Jack D. Adderley, Simona John von Freyend, Sarah A. Jackson, Megan J. Bird, Amy L. Burns, Burcu Anar, Tom Metcalf, Jean-Philippe Semblat, Oliver Billker, Danny W. Wilson, Christian Doerig

## Abstract

Intracellular pathogens are known to mobilise host signaling pathways to manipulate gene expression in their host cell to promote their own survival. Surprisingly, evidence is emerging that specific signal transduction elements are activated in a-nucleated erythrocytes in response to infection with malaria parasites, but the extent of this phenomenon remains unknown. Here, we fill this knowledge gap by providing a comprehensive and dynamic assessment of host erythrocyte signaling during the course of infection with *Plasmodium falciparum*. We used an antibody microarray that comprises 878 antibodies directed against human signaling proteins, >600 of which are phospho-specific, to interrogate the status of host erythrocyte signaling pathways at three stages of parasite development during the asexual cycle. This confirmed the pre-existing fragmentary data on the activation of a PAK-MEK pathway, and revealed the modulation of a large number of additional signaling elements during infection.

We focussed on the receptor tyrosine kinase c-MET, also known as the hepatocyte growth factor receptor, and the MAP kinase pathway component B-Raf that is reported to lie downstream of c-MET in a number of cell types. Array data validated by Western blotting revealed that activation sites of c-MET are phosphorylated in trophozoite-infected erythrocytes, and we show that treatment of parasite cultures with c-MET or B-Raf selective inhibitors have nanomolar potency against *in vitro* proliferation of *P. falciparum* and the phylogenetically distant species *P. knowlesi*. Furthermore, we demonstrate that a c-MET inhibitor impairs *in vivo* proliferation of the rodent malaria parasite *P. berghei* in mice.

Overall, we provide a comprehensive dataset on the modulation of host erythrocyte signaling during infection with malaria parasites, as well as a proof of concept that human signaling kinases identified as activated by malaria infection represent attractive targets for antimalarial intervention.

## Introduction

Malaria, caused by infection of mosquito-borne apicomplexan parasites in the genus *Plasmodium*, remains one of the most devastating infectious diseases globally. Among the six species that infect humans, *P. falciparum* is the most virulent and is responsible for the majority of malaria-related deaths. The remaining human malaria species *P. vivax, P. ovale, P. malariae* and two zoonotic species, *P. knowlesi* and *P. simium* ^1,2^, are major contributors to global malaria morbidity and must be considered in the context of new treatment strategies for malaria. Recent years have seen a 50% drop in malaria-related mortality, yet the disease still kills an estimated 445,000 people every year, mostly young children in sub-Saharan Africa ^3^. Progress towards malaria elimination and ultimately eradication has stagnated, and the emergence of parasites resistant to the most recently deployed global front-line treatment, artemisinin-combination therapies (ACT) ^3,4^, is a major concern. It remains critical to develop novel antimalarial drugs with untapped modes of action.

Protein kinases (PKs) are core components of signaling pathways in eukaryotic cells. Phosphorylation of the target protein can lead to conformational changes and generate or mask binding motifs, and thus can affect its activity, binding properties, stability or subcellular localisation. PK activity can be regulated positively or negatively by phosphorylation of specific amino acids, or by binding to specific activator or inhibitor proteins. Activation leads to a conformational change making the active site accessible to ATP and the substrate protein. PKs are eminently druggable targets, as illustrated by the fact that > 48 small molecule kinase inhibitors have reached the market, mostly in the context of cancer chemotherapy ^5,6^. The kinomes of apicomplexan parasites and humans are quite divergent owing to the large phylogenetic distance between these organisms ^7^, and parasite-encoded PKs have been proposed as attractive potential targets for selective intervention ^8^. Host-directed therapy (HDT) is another avenue that is currently gaining traction to combat infectious diseases generally ^9^. We and others ^10,11^ have proposed that host erythrocyte (and hepatocyte) PKs represent excellent potential targets for antimalarial intervention. Many anti-infective drugs are rendered ineffective by the selection of mutations in their pathogen-encoded targets, which explains the rapid emergence of resistance across essentially all pathogen taxons. Targeting host enzymes would deprive the pathogen of this most direct pathway to resistance. Furthermore, repurposing anti-cancer kinase inhibitors as agents against infectious diseases would greatly alleviate the lack of resources that severely affects anti-infective drug development.

Malaria pathology is caused by the blood stage of the parasite lifecycle. Blood stage malaria begins with the infection of erythrocytes by *Plasmodium* merozoites, followed by rapid growth and asexual multiplication. Newly formed merozoites are then released into the bloodstream and invade erythrocytes to begin the next replication cycle (~48 hrs for *P. falciparum*). Several human signaling molecules of the host erythrocyte have been implicated in parasite survival during infection. These include G-coupled protein receptors ^12–14^, the protein kinases MEK (MAP/ERK kinase) ^15^, PAK (p21-activated kinase) ^15^, Protein kinase C (PKC) ^16^, and peroxiredoxin ^17–19^. In our study of the erythrocyte PAK-MEK pathway ^15^, we showed that host MEK1 is phosphorylated on residue Ser298 in infected erythrocytes. pSer298 is known to cause activation of the enzyme through stimulating trans-auto-phosphorylation on its activation loop. The only PK known to phosphorylate MEK1 on Ser298 is PAK1 ^20^, and indeed we showed that the latter enzyme is also activated in infected erythrocytes ^15^. PAK isoforms are notorious for having a multitude of substrates ^21^ and serve as a node to integrate signals from a number of upstream receptors and transmit these signals to several downstream effector pathways (including the mitogen-activated protein kinase pathway [MAPK] that includes the aforementioned MEK). We, therefore, hypothesised that our description of the PAK-MEK pathway addresses but a minute fraction of the signaling events that may be modulated by infection. Here, in order to comprehensively assess the host erythrocyte signaling response to infection, we use a microarray of antibodies designed to recognise human signaling proteins and their phosphorylation status. Comparison of the signals obtained from extracts of uninfected erythrocytes with those obtained from erythrocytes infected with *P. falciparum* at three main developmental stages of the replication cycle (rings, trophozoites and schizonts) allowed us to generate a comprehensive and dynamic picture of the modulation of host erythrocyte signaling by infection. This identified several host kinases as potential targets for HDT; on this basis, we further demonstrate that selective inhibitors against human c-MET and B-Raf display high potency against *P. falciparum* and *P. knowlesi in vitro*, and show that a c-MET inhibitor has *in vivo* activity against *P. berghei* in a murine model of malaria.

## Results

### Kinexus antibody microarray analysis

To investigate dynamic changes in host erythrocyte signaling during *P. falciparum* asexual proliferation, we employed an antibody microarray developed by Kinexus (Vancouver, Canada). The array consists of 878 unique antibodies, 265 of which are pan-specific, i.e. recognise both the phosphorylated and unphosphorylated forms of the target protein; the remaining 613 antibodies are phosphorylation-specific, recognising signaling molecules only if their activating or inhibitory phosphorylation sites are modified by the addition of a phosphate group. Some of the most important and well-known signaling molecules, such as members of the mitogen-activated protein kinase (MAPK) pathways or the PKC isoforms, are detected by multiple antibodies aimed at various phosphorylation sites within the same protein. The array thus provides a comprehensive picture as to how signaling mediated by these molecules changes during infection. Each array device comprises two identical chambers, each carrying two spots for each of the 878 antibodies, thus delivering each read-out in duplicate. Two sample lysates are labelled with protein-binding fluorescent dye [e.g. from *Plasmodium*-infected (iRBC) and non-infected (uRBC) red blood cells] and compared on each device, one sample per chamber. Binding of proteins to the antibodies on the array is quantitated through signal intensity for each spot.

### Robustness of the approach

To evaluate the reproducibility and robustness of the approach, we compared four uninfected erythrocyte (uRBC) samples from different blood donors that were cultured for ~24 hrs under the same conditions as the iRBCs. The mean signal intensity and standard deviation for each phospho-specific antibody on the array were determined for these four samples (Fig. 1; the entire datasets are available in Supplemental dataset 1). Despite some variation, individual data points for each antibody remained within two standard deviations of their mean, illustrating that sample-to-sample variation was minimal.

**Figure 1.**
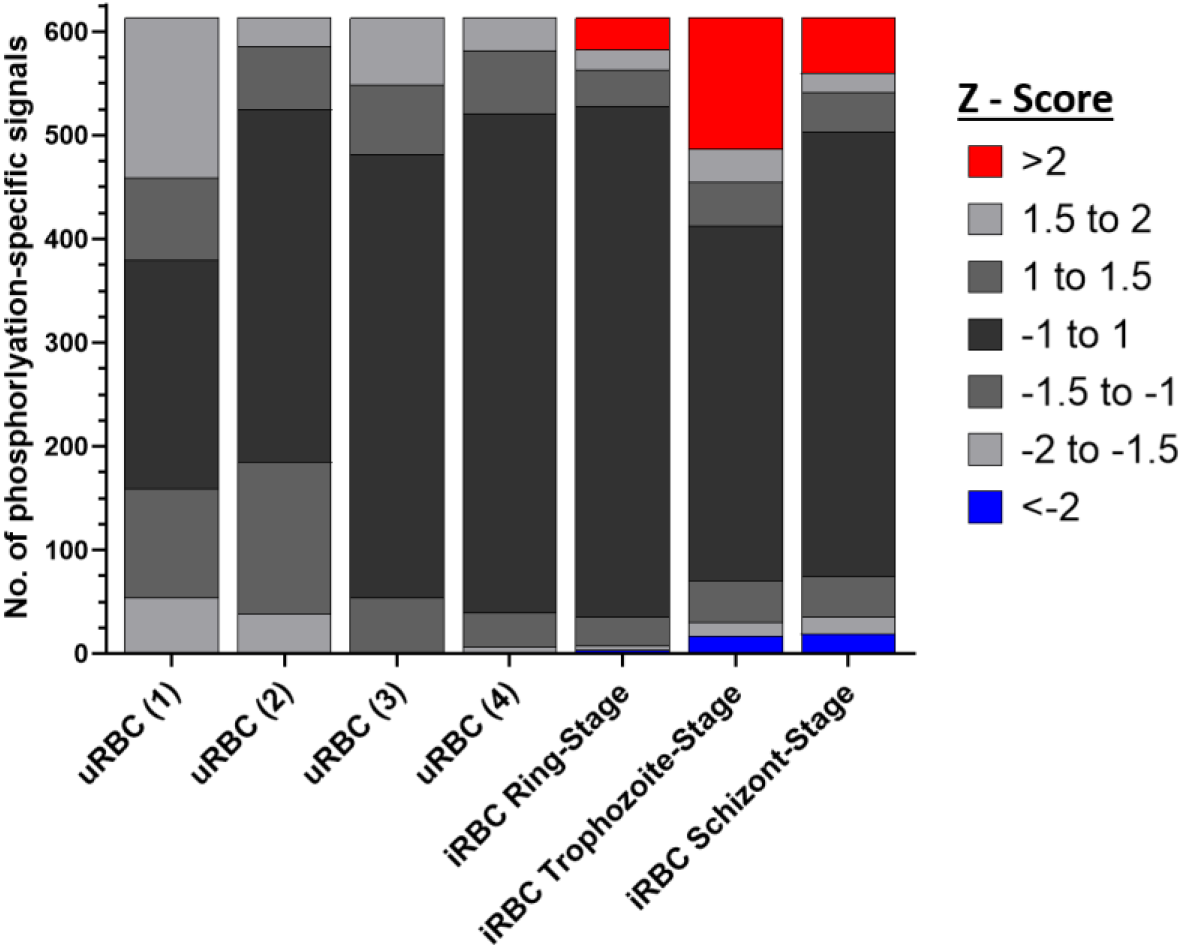
Distribution of Z-scores for each sample. All antibody microarray signals from each sample (four from uRBCs and the mean signal for each iRBC harboring parasites at the ring (n=3), trophozoite (n=3) and schizont stage (n=2)) were compared to the uninfected erythrocyte mean value for each signal, and the Z-score determined for each phosphorylation-specific signal. The Z-score is defined as the number of standard deviations from the uRBC mean for each sample; a positive or negative value indicates the direction of the signal (increase or decrease, respectively) from the uRBC mean. No signals from the uRBC samples (n=4) had associated Z-scores above 2 or below −2, while the three iRBC time points showed clear variations from uRBCs.

### Overall changes in phosphosignaling in response to infection

To assess the impact of infection on host cell signaling, whole-cell protein extracts were obtained from synchronised *P. falciparum* cultures at three time windows during the erythrocytic cycle: 4-12 <u>hr</u>s <u>p</u>ost-<u>i</u>nvasion (hpi) (rings, n=3), 24-28 hpi (trophozoites, n=3) and 44-48 hpi (segmented schizonts, n=2). Each of these extracts were loaded onto a separate antibody array and compared to the mean of the uRBC lysates harvested in parallel and containing RBCs from the same donor as the iRBC sample (see above). In stark contrast to the minimal variation between the uRBC samples (see above), analysis of the iRBC extracts showed that 6% (ring-stage), 23% (trophozoite-stage) and 12% (schizont stage) of the phospho-specific antibody intensities differed by more than two standard deviations from uRBC mean. The datasets for each of the three developmental stages showed no internal variation greater than two standard deviations between the biological replicates (Supplemental Figure 1), further confirming the robustness and reliability of the approach (Fig. 1; the complete datasets are available in Supplemental Dataset 1). Overall, this implies (i) that there is little variation between biological replicates of uRBCs, attesting to the reliability of the system, and (ii) that infection with *P. falciparum* causes significant variation in the signals yielded by phospho-specific antibodies, suggesting that *P. falciparum* strongly impacts host erythrocyte phosphosignalling during infection.

### Data filtering-removal of cross-reacting antibodies

To address possible cross-reactivity of the antibodies with parasite-derived proteins, we compared signals from purified unsynchronised parasites (pellet obtained by saponin lysis followed by centrifugation) with those of the purified erythrocyte cytoplasm (saponin supernatant). Saponin disrupts the erythrocyte membrane, thereby releasing erythrocyte cytoplasmic proteins and exported parasite proteins, while the parasite and insoluble erythrocytic material can be pelleted ^22^. The array was loaded with 20x more (protein mass) pellet material than supernatant material, to confer high stringency to the cross-reactivity filter. A heatmap of the results is shown in Supplemental Figure 2a (full data available in Supplemental Dataset 2). Antibodies showing a fold change > 1.5 between the erythrocyte cytoplasm (saponin supernatant) and the parasite extract (saponin pellet), amounting to 224 signals (37%) of the phospho-specific antibodies, were withdrawn from further analysis (Supplemental Figure 2 and Supplemental Dataset 2). This level of cross-reactivity is not surprising, as many signaling proteins display conservation between *Homo sapiens* and *P. falciparum*, especially in their regulatory sites ^7^. Some of these cross-reactive antibodies may recognise parasite-encoded orthologues of the human target proteins and thus prove to be useful tools to study *P. falciparum* signal transduction; however, this lies outside of the scope of the present study.

### Data filtering - removal of low-signal antibodies

Some antibodies displayed a weak fluorescence signal, likely due to the low abundance of the target protein. The antibodies yielding a signal intensity below a fluorescence reading of 1000 units in both the erythrocyte control and parasite-infected samples were removed from further analysis, as recommended by Kinexus (Supplemental Dataset 1). This included 26 signals from the ring array, and 41 and 48 signals from the trophozoite and schizont arrays, respectively.

### Broad analysis of the post-filtering iRBC dataset

Following low and cross-reactive signal removal, 1 of the ring stage signals, 29 of the trophozoite stage and 17 of the schizont stage signals had fold changes greater than 2 or below 0.5 compared to their uninfected counterpart, revealing dynamic changes in the phosphorylation of host signaling proteins during *P. falciparum* asexual development (Fig. 2a: heatmap of all retained signals; Fig 2b: distribution of increased and decreased signals at the three development stages). The small number of changes in ring-infected cells may in part be due to the fact that these samples contained only 33% infected cells (as magnetic purification of infected cells, which allows close to 100% iRBCs for the trophozoite and schizont stages, cannot be implemented for ring stages). The observed effect on host cell phosphosignaling was considerably larger at later stages of infection (trophozoite/schizont), with the majority of these changes being attributed to increases in phosphorylation (80% and 65%, respectively). A dot plot for each of the three life stages (Fig. 2c) further illustrates this trend and highlights the strongest individual changes.

**Figure 2.**
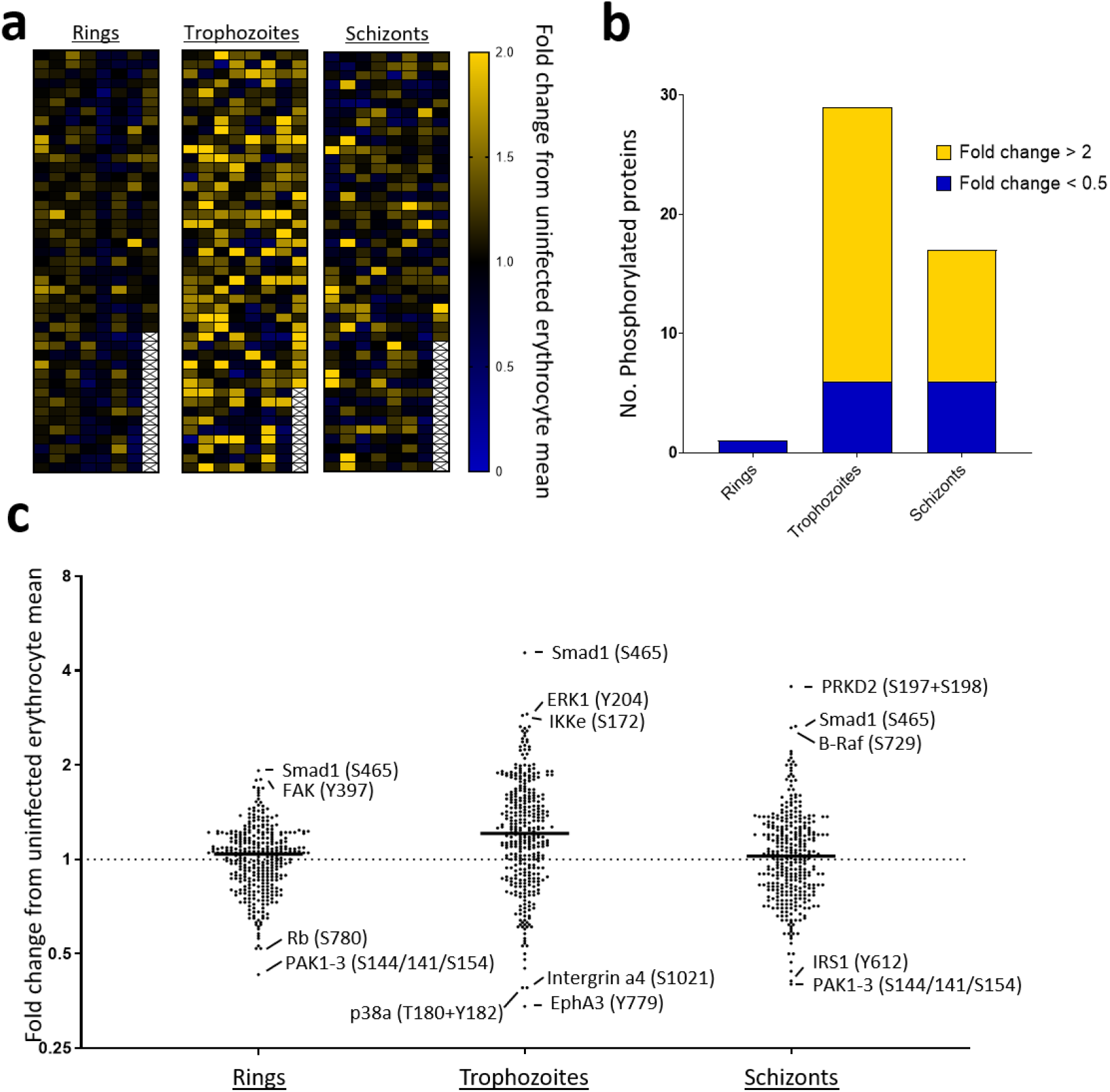
Microarray data for the ring, trophozoite and schizont stages. **a)** Heat map of phosphospecific antibody array data for ring-trophozoite- and schizont-infected cells after removal of cross-reactive and weak signals. **b)** Phosphorylation-specific signals (post-filtering) ranked according to fold change. Yellow represents changes with a fold change greater than 2; blue represents changes with a fold change less than 0.5. Full datasets are presented in Supplemental Dataset 1 and 2. **c)** Dot plot highlighting differences in phosphosite signals represented as fold change. Strong changes are listed with their associated phosphorylation site.

### Shortlisting of high-confidence changes in phosphosignaling

Of the reliable antibodies remaining after removal of low and cross-reactive signals, 16/345 (ring stage), 102/351 (trophozoite stage) and 35/346 (schizont stage) were significantly different (p<0.5, unpaired t-test) from the uninfected erythrocyte signals. Figure 3 lists the most noteworthy changes observed from this analysis (the full list of the significant changes is available in Supplemental Dataset 3).

**Figure 3.**
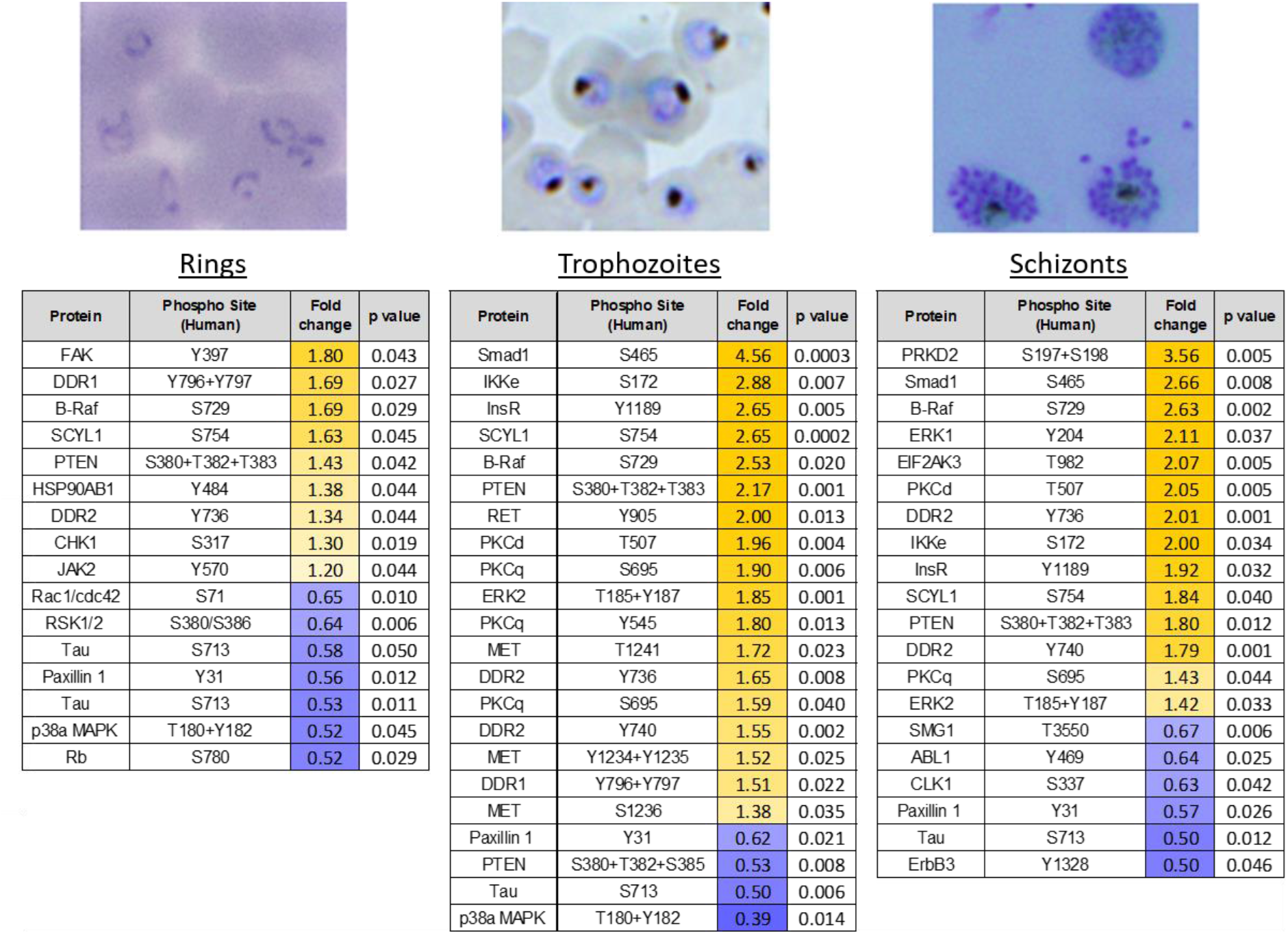
Shortlist of the highest confidence signals (p<0.05) for rings, trophozoites and schizonts. Increases in phosphorylation compared to uninfected erythrocytes are marked in yellow; decreases are marked in blue. All values listed are the fold change from the uRBC control mean (n=4); the corresponding p-value for each signal is also listed. Each signal has a p-value <0.05, as determined through an unpaired t-test. All listed signals passed stringent parasite cross-reactivity and signal intensity filters (see text for details and Figure 2).

Some of the most notable changes implicated the PKC isoforms δ and θ. Several PKC isoforms have previously been reported to be activated during *P. falciparum* infection of erythrocytes, but the phosphorylation status of these kinases was uncharacterised^23,24^. Several phosphorylation sites of PKCδ/θ were significantly phosphorylated in trophozoite- and schizont-infected cells (Fig. 3; a list of the phosphorylation specific signals for PKCδ/θ is available in Supplemental Figure 3). The most striking changes were observed for Thr507 (PKCδ) and Ser695 (PKCθ). Thr507 is a regulatory residue located within the activation loop of PKCδ^25^ while, Ser695 is within the hydrophobic motif of PKCθ, and its phosphorylation is required for optimal kinase activity^26^. Additionally, phosphorylation of Ser695 may also be involved in the localisation of PKCθ to the plasma membrane ^27^. These data are consistent with previous reports that host cell PKC activity is detectable during schizont development (30-40 hrs post-invasion) with limited activity before 25 hrs ^23,24^, and considerably refine these previous findings, notably with respect to dynamic phosphorylation changes on specific residues of the various PKC isoforms during parasite development.

### MAP-Kinase signaling in infected erythrocytes – B-Raf, MEK and ERK

The canonical MAPK pathway is a highly conserved, three-tiered cascade, in which a MAP kinase kinase kinase (MAP3K, or MEKK for MAP/ERK kinase kinase) phosphorylates and thereby activates a MAP2K (or MEK), which in turn phosphorylates and activates a MAP kinase. A well-understood MAPK pathway involves Raf, MEK and ERK. Both Raf isoforms are activated by Ras-GTPase binding, which enables phosphorylation of T598 and S601 (on B-Raf) and S338 and Y340 (on C-Raf) ^28–30^. This results in the full activation of Raf, enabling downstream phosphorylation of MEK and consequently ERK1/2 (pathway depicted in Fig. 4b). *P. falciparum* has no homologues for either Ras, Raf or MEK, but expresses two MAPKs, neither of which is a close homologue of ERK1/2 ^31^. Figure 4 lists the phosphorylation sites detected on the array that are relevant to canonical MAPK pathways (Fig. 4a) and indicates the activating phosphorylation sites for each of the three kinases involved (Fig. 4b).

**Figure 4.**
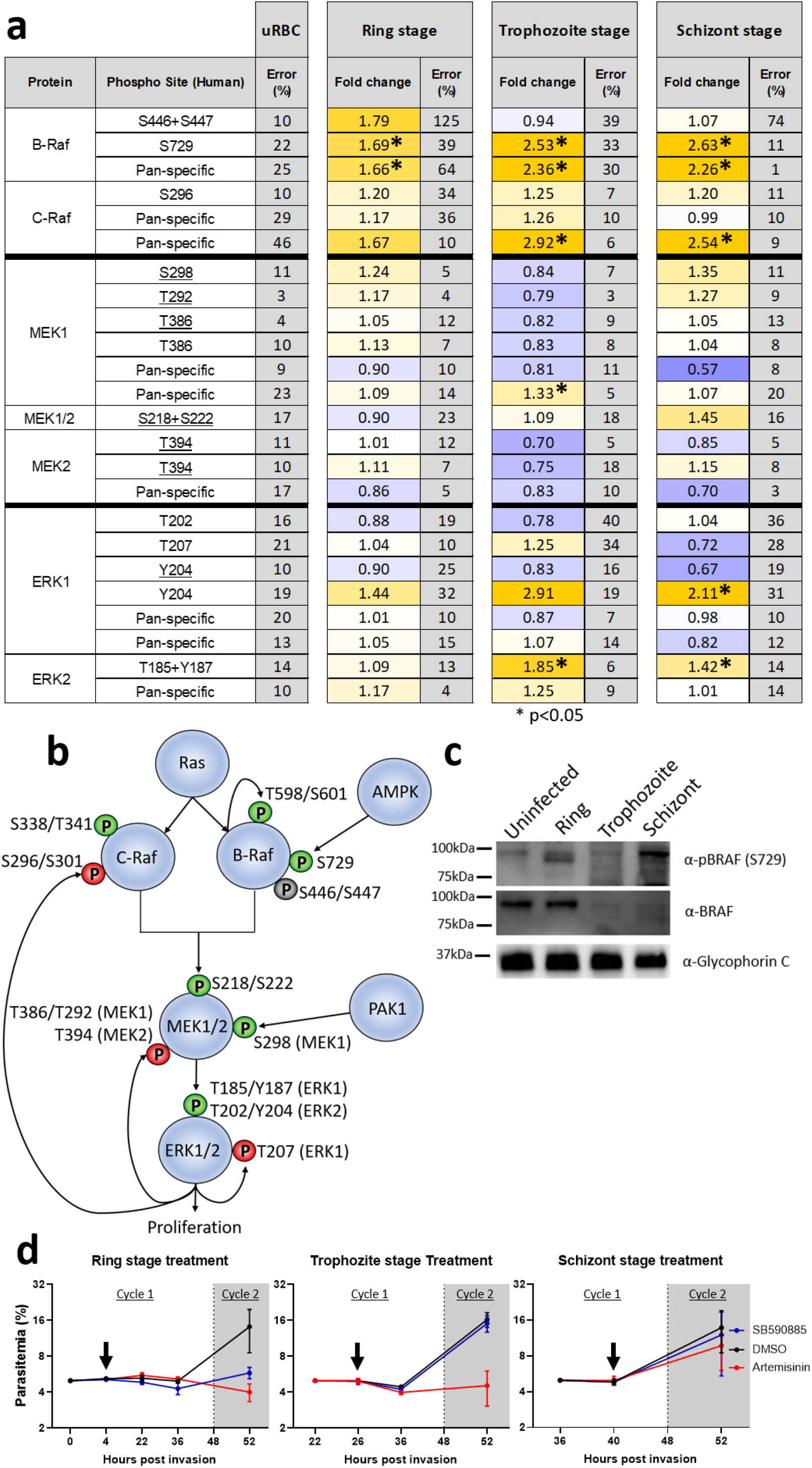
Activation of a MAPK pathway in infected erythrocytes. **a)** Summary table of the MAPK pathway components and phosphorylation sites covered by the antibody microarray, including RAF, MEK1/2 and ERK1/2 with the fold change from the uninfected erythrocyte control signal for each of the three time points (erythrocytes infected with rings, trophozoites or schizonts) shown. Signals which were significantly different during infection are marked by * (p<0.05, unpaired t-test). Phosphosites underlined in the table were flagged as cross-reactive with parasite material in Fig. 2, but are included here for completeness. **b)** Schematic representation of the MAPK pathway with the relevant phosphorylation sites present in the array data set covered in **a**. Green ‘P’ represents phosphorylation sites associated with the activity of the respective kinase, while the red ‘P’s represent negative feedback phosphorylation sites. The grey ‘P’ represents a known priming site on B-Raf. **c)** Western blot validation of B-Raf phosphorylation. A phospho-specific antibody to B-Raf (S729) detected phosphorylated B-Raf in ring-stage infected erythrocytes (upper panel). A pan-B-Raf antibody detected B-Raf in the uninfected and ring-stage infected erythrocyte samples, with a strong decrease at the trophozoite and schizont stages (middle panel). An antibody against Glycophorin-C was used as a loading control (lower panel). **d)** Synchronised *P. falciparum* cultures were treated in 4-hr windows with 2.45 μM SB590885 (5x IC_50_, see Fig 6), 0.091 μM artemisinin or DMSO (vehicle control), starting at 0 (rings), 22 (trophozoites) and 38 (schizonts) hours post-invasion (the black arrows indicate the end of treatment, where the medium was replaced with fresh medium without drug). The parasitemia was then monitored until the next cycle of parasite development to assess parasite survival. Starting parasitemia was normalised to 5% across the biological replicates (n=3) error bars represent the standard error of the mean.

Proteins in the MAPK signaling pathway showed notable changes in phosphorylation in iRBCs compared to uRBCs. B-Raf and C-Raf as the ‘gatekeeper’ proteins of this pathway showed changes in phosphorylation patterns with *P. falciparum* RBC infection. Unfortunately, the activating Ras-induced phosphorylation sites on B-Raf (T598 and S601) and C-Raf (S338 and Y340) are not represented in the array and could not be assessed. Antibodies against C-Raf S296 yielded a consistent fold change of >1.2 for all three parasite stages. Phosphorylation of C-Raf on S296 is thought to be mediated by ERK1/2 in the context of a negative feedback loop ^32^, which would be consistent with activation of the MAPK pathway detected here (see below).

B-Raf phosphorylation at S446 and S447, sites associated with priming B-Raf for activation by Ras ^30,34^, increased by 1.79-fold at the ring stages, but decreased back to uRBC levels in trophozoites and schizonts. B-Raf S446 is constitutively phosphorylated in some cell lines, but this remains to be shown in cells of the erythropoietic lineage ^30,34–36^. In contrast, phosphorylation of B-Raf at S729, a Ras-independent phosphorylation event carried out by AMP-activated protein kinase (AMPK) in response to energy stress, increased at all three life cycle stages with a fold change ranging from 1.69 to 2.63 across parasite development (Fig 3, Fig. 4a). Of the two AMPK antibodies on the array, one was discounted due to cross-reactivity with a parasite protein, while the other (AMPKa2 S377) shows a slight increase (1.17 fold change) in phosphorylation at the trophozoite time point supporting that this kinase may also be activated in iRBCs (Supplemental Dataset 1).

We next determined the total amount of B-Raf and phosphorylated B-Raf S729 at each lifecycle stage using Western blot. Detection of total B-Raf protein using a pan-antibody independent of phosphorylation showed that the amount of full-length B-Raf was greatly reduced at trophozoite and schizont stages compared to uRBCs and rings (Fig. 4c). This fits with the reduced phosphorylation identified for S446 and S447 at trophozoites and schizonts stages. Paradoxically, while we observed increasing S729 and total B-Raf signals from rings to schizont stages on the array, this was not reflected in increased signal for full-length B-Raf on the Western Blot. Instead, the increasing signal on the array corresponded to increasing bands on the Western blot that have lower molecular mass (see Supplemental Figure 4 for additional data). Together this indicates that B-Raf is indeed phosphorylated at the ring stage of *P. falciparum* development, before degradation in the later stages of parasite development (see Discussion).

Overall, the microarray and Western blot data points to activation of B-Raf at ring-stages, and degradation at later stages. To assess if parasite requirements for host B-Raf activation were also ring-stage-specific, as suggested by the apparent degradation of B-Raf in trophozoite- and schizont-infected cells (Fig. 4c), we treated highly synchronized parasites in 4-hour windows with a potent B-Raf inhibitor (SB-590885, see below). Only when the treatment was applied at ring-stage did it display parasiticidal activity, whereas there was no effect when the cultures were treated at the trophozoite or schizont stage, in stark contrast to artemisinin, which displayed (as expected) a strong effect at the trophozoite stage (Fig. 4d).

Further supporting activation of Rafs in iRBCs, the B-Raf/C-Raf substrates MEK1/2 showed increased phosphorylation on several residues at both the ring and schizont time points (Fig. 4a). MEK1/2 phosphorylation of S298 (resulting from p21-activated kinase [PAK] activity) during the late stages of *P. falciparum* infection has previously been established by Western blot analysis ^15^, aligning with the 1.35-fold increase of pS298 at the schizont stage observed in the array experiment. Other phospho-sites such as T292 and S218+S222 (MEK1/2) show a signal increase in schizont-infected red cells. MEK1 T292 is involved in negative feedback of the kinase ^37^, while S218+S222 and S222+S226 are the activation loop phosphosites in MEK1 and MEK2, respectively ^38,39^. All the MEK1/2 phospho-specific antibodies, as well as the ERK1 (Y204) antibody, showed some cross-reactivity with parasite protein (underlined in Fig. 4a), which makes the overall MEK1/2 data difficult to interpret (Fig. 4a).

MEK1/2 phosphorylate ERK1 and ERK2 (on T202-204 for ERK1, on T185-Y187 for ERK2), and these residues are the only known targets of MEK1 activity ^40,41^. ERK2 is slightly more phosphorylated in ring-infected cells than in uninfected erythrocytes (fold change = 1.09); at the trophozoite stage it has increased further (fold change = 1.85), and at the schizont stage ERK2 phosphorylation is still elevated (fold change = 1.42). The array results for the ERK1 activation sites (T202 and Y204) at the ring-stage of *P. falciparum* development are however difficult to interpret as these signals indicate increases as well as decreases. ERK1 T207 is a known autophosphorylation regulatory site and showed a fold change value of 1.25 in trophozoite-infected red cells-, supporting that regulation of ERK1 activity occurs during infection ^42^. These data support that *P. falciparum* iRBCs modify the host cell Raf/Mek/ERK pathway during growth and that the activity of RBC B-Raf is required for parasite maturation beyond ring stages.

### c-MET signaling in infected erythrocytes

Following the identification of the MAPK pathway activation, we focused our attention on signaling components that lay upstream of the MAPK pathway and found that the receptor tyrosine kinase c-MET, which is known to signal to MAPK pathways in other systems ^43^, was clearly activated by infection (Fig. 3). C-MET is expressed in many cells of mesenchymal origins ^44,45^, and its activation occurs through the binding of hepatocyte growth factor (HGF) to the extracellular domain, prompting homodimerization and trans-autophosphorylation and activation ^46^. C-MET controls several key downstream pathways including (but not limited to) the MAPK pathway (proliferation, motility and cell cycle progression) and the phosphatidylinositol 3-kinase (PI3K)/AKT pathway (cell survival) ^43^. There is no homologue of c-MET (or any member of the Tyrosine kinase group) in *P. falciparum* ^7^.

The array contained a total of nine phospho-specific antibodies targeting c-MET, 7 of which passed cross-reactivity and low-intensity signal thresholds (Fig. 5a). There was no noteworthy change in the phosphorylation of any of these residues at the ring stage. In contrast, five out of the seven sites show an increase in phosphorylation in trophozoite-infected cells (fold change >1.38, Fig. 5a). The signals from two antibodies against c-Met phosphorylation sites decreased at the trophozoite stage. One of these is directed against Y1230+Y1234+Y1235; this antibody is affected by high error between duplicates (848% at the trophozoite stage) and was deemed unreliable. The other antibody recognises pY1230 and is the only reliable c-MET antibody to show a decrease of phosphorylation at the trophozoite stage. At the schizont stage, most sites were slightly higher than the uninfected control. Two unique pan-specific c-MET antibodies were also present on the array. Both pan-specific antibodies showed minimal signal variation between samples, indicating that the level of c-MET remained relatively constant as expected (Fig 5a).

**Figure 5.**
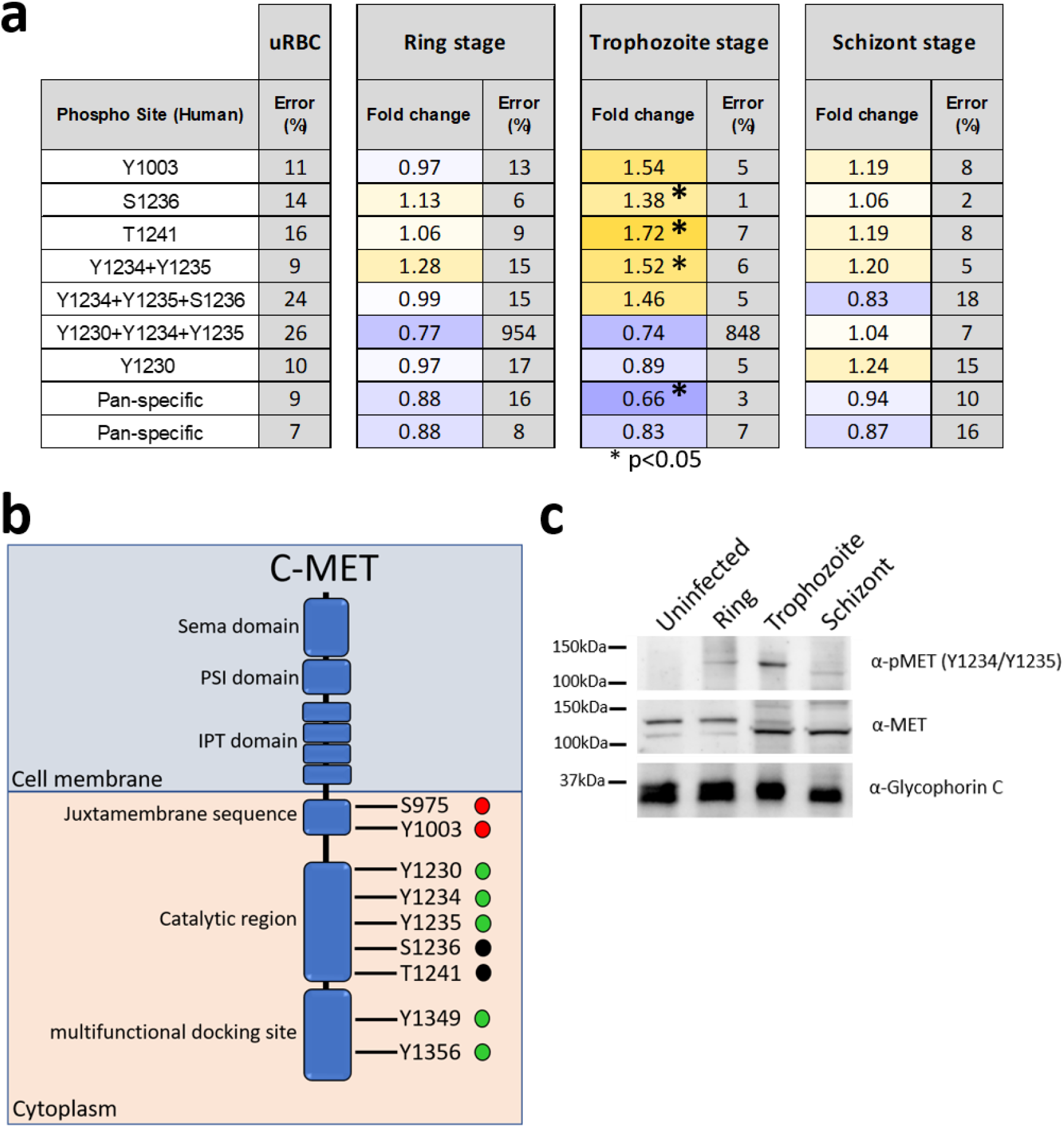
Activation of c-MET in infected erythrocytes. **a)** c-MET phosphorylation sites covered by the antibody microarray, with the fold change from uRBC control values for the three-time points (ring, trophozoite and schizont). Signals which were significantly different during infection are marked by * (p<0.05, unpaired t-test). **b)** Schematic of the c-MET protein. The juxtamembrane sequence contains both S975 and Y1003, which represent c-CBL binding sites that target the receptor for degradation (red circles) ^51,52^. The catalytic domain contains Y1230, Y1234, Y1235, S1236 and T1241. All activating tyrosine phosphorylations (green circles) occur through receptor homodimerisation ^43,46,53^. The function of S1236 and T1241 (black circles) is uncharacterised. The multifunctional docking site contains two phosphosites, Y1349 and Y1356, which aid in the binding of adapter molecules such as grb2 and GAB1 ^46^. **c)** Western blot validation of c-MET phosphorylation across asexual development. A phospho-specific antibody to c-MET (Y1234+Y1235) detected phosphorylated c-MET at the ring and trophozoite stages (upper panel). A pan c-MET antibody detected c-MET across all four samples, with a double band in the uninfected, ring and trophozoite samples (middle panel). An antibody against Glycophorin-C was used as a loading control (lower panel).

Antibodies against c-MET S975 and Y1349+Y1356 were not present on the array or removed from analysis because of low-intensity signal, respectively. Reliable antibodies were available for pY1230 and pY1234+pY1235, which lie within the c-MET catalytic domain and are phosphorylated in response to binding of HGF, activating the kinase^47^. While increased phosphorylation was evident for Y1234+Y1235 across lifecycle stages, with a peak signal seen at trophozoites, this occurred only at schizont stage for Y1230. Increased Y1234+Y1235 phosphorylation compared to uRBCs was confirmed by Western blot analysis using independently-sourced Y1234+Y1235 and pan-MET antibodies (not used on the microarray), with peak signal again seen in trophozoites (Fig. 5c). This confirms that c-MET is indeed phosphorylated in the catalytic domain during trophozoite infection. As observed for B-Raf, we saw a decrease in apparent size of c-MET during the ring-to trophozoite transition, which may be due to differences in phosphorylation status, or to partial degradation of the protein (see Discussion).

Phosphorylation of c-MET Y1003 peaks at the trophozoite stage. In nucleated cells, phosphorylated Y1003 recruits the c-CBL (Casitas B-lineage lymphoma) protein, which promotes ubiquitination and degradation of c-MET via the 26S proteasome ^48,49^. The activity of the 20S proteasome has been previously reported in mature erythrocytes; in contrast, no 26S activity was observed in these cells ^50^. The absence of 26S proteasomal activity in erythrocytes could explain why c-MET protein maintains a steady-state level throughout blood-stage parasite development (Fig. 5a), despite Y1003 phosphorylation.

### Small-molecule inhibitors of host kinases identified with the array impair parasite proliferation

We previously published that U0126, a highly selective allosteric inhibitor of MEK1/2, as well as other structurally distinct selective inhibitors of these kinases, display low μM potency against *P. falciparum in vitro* ^15^. The findings reported here that several additional human kinases such as c-MET and B-Raf are activated by infection raise the question of whether their inhibition would likewise impair parasite proliferation. Several selective inhibitors of c-MET and B-Raf have emerged from the cancer drug discovery pipeline, allowing us to measure the IC_50_ values in parasite proliferation assays of the following inhibitors: Crizotinib (an inhibitor of c-Met and the related receptor tyrosine kinase ALK [anaplastic lymphoma kinase]); PHA-665752 (a c-MET-selective inhibitor), SB-590885 (a B-Raf selective inhibitor) and Sorafenib (a B-Raf inhibitor which also targets C-Raf, albeit with a 10-fold lower potency) ^54^ (Fig. 6b&c). Despite the absence of B-Raf, c-MET and ALK homologues in the *Plasmodium* kinome, all these inhibitors displayed low nano-to micromolar IC_50_ values against both *P. falciparum* and *P. knowlesi* in parasite proliferation assays (Fig. 6a). We cannot at this stage exclude off-target effects against *Plasmodium* proteins. However, our observation that the B-Raf inhibitor SB-590885 is effective only against ring stages, i.e. during the period where B-Raf is present and before its apparent degradation during the ring to trophozoite transition (Fig 4a&d), together with the absence of c-MET or B-Raf homologues in the parasite’s kinome, strongly suggests that inhibition of these host erythrocyte kinases impairs parasite proliferation. This in turn indicates that activation of these human kinases by malaria parasites may play an important role in their survival inside erythrocytes.

**Figure 6.**
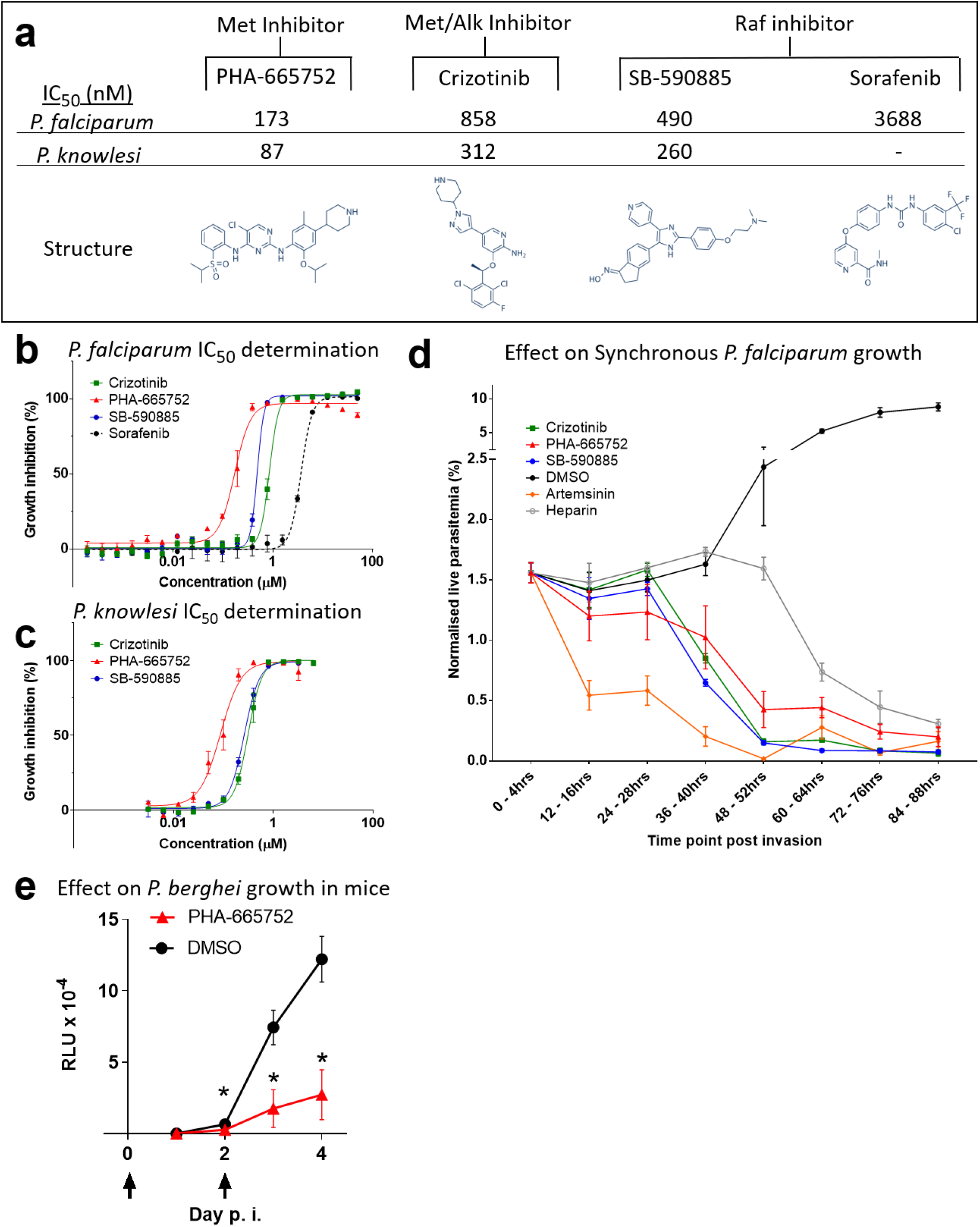
IC_50_ and phenotypic effect of human kinase inhibitors on *P. falciparum* and *P. knowlesi* blood-stage development. **a)** IC_50_ values of the human kinase inhibitors Crizotinib (MET/ALK inhibitor), PHA-665752 (Met inhibitor), SB-590885 (B-Raf inhibitor) and Sorafenib (B-Raf inhibitor, *P. falciparum* only) on asynchronous *P. falciparum* and ring-stage *P. knowlesi* asexual growth. **b)** Growth inhibition curve for asynchronous *P. falciparum*, IC_50_ measured after 72 hrs of compound exposure, n=3, n=2 for sorafenib. **c)** Growth inhibition curve for *P. knowlesi*, IC_50_s measured after 50 hrs of compound exposure, n=3. **d)** Effect of treatment with 5x the IC_50_ value of the compounds Crizotinib, PHA-665752, SB-590885, Artemisinin, DMSO (equivalent volume of DMSO in Crizotinib sample) and heparin (15 μl/ml) on highly synchronous (4-hr window) *P. falciparum* cultures over 88 hrs (n=5). The Y-axis represents the live parasitemia values for each time point normalised to a starting total parasitemia of 2%. All error bars represent the standard error of the mean. **e)** Effect of treating mice with PHA-665752 on the growth of *P. berghei*. The graph shows relative luminescence units (RLU) in the blood of BALB/c mice infected with parasites constitutively expressing firefly luciferase and treated or not with PHA-665752 at 25 mg/kg at times indicated by arrows. * = p < 0.001 by two-tailed *T*-test; n = 7 mice per treatment group.

To determine at which point in the asexual cycle the parasite die if treated with c-MET inhibition at the start of the cycle, and to expand the aforementioned data pertaining to B-Raf inhibition (Fig. 4c), cultures of *P. falciparum* were highly synchronised to a 0 – 4 hrs post-invasion window of development. Cultures were treated with 5x IC_50_ of SB-590885, Crizotinib, PHA-665752, the frontline antimalarial Artemisinin, or with the merozoite invasion inhibitor heparin. Parasites were monitored via flow cytometry every 12 hr over 84 hrs. The normalised live parasitemia detected for each time point was determined through co-staining with Hoechst 33342 (DNA stain) and Mito-tracker Orange (mitochondrial potential stain). The average values of the biological replicates are represented graphically in Fig. 6d. Treatment with Artemisinin resulted in a reduction in live parasitemia at 12-16hpi, indicative of ring-stage parasite death. Treatment with heparin caused a drop in live parasitemia post 48hpi, as expected for an invasion inhibitor. In contrast, the first notable reduction in parasitemia observed after treatment with Crizotinib, PHA-665752 or SB-590885 occurs prior to the schizont stage. This indicates that all three human kinase inhibitors exert their largest effect on parasite survival at a time that is consistent with the activation time of their target kinases (and, in the case of B-Raf, with the time where the full-length protein is present, i.e. ring stage), as revealed by the microarray and Western blot analysis. Notably, all compounds except heparin resulted in a live parasitemia below 0.2% by 84-88hpi, with SB-590885 and Crizotinib reducing live parasitemia to <0.1%. For an initial assessment of the *in vivo* efficacy of the c-MET inhibitor PHA-665752, we treated mice infected with the rodent malaria parasite *Plasmodium berghei*. A treatment regime and formulation that was previously used to control tumour growth in mice with the same inhibitor ^56^ led to a 3.3-fold reduction in parasite load after two doses of treatment *in vivo* (Fig. 6e, day 4).

## Discussion

The antibody microarray analysis generated a large number of testable hypotheses on the involvement of host erythrocyte signaling proteins during infection by the human malaria parasite *P. falciparum*. We achieved Western blot validation for three of the host kinases (MEK1 [15], c-Met [this study] and B-Raf [this study]) that the array data suggest are activated by infection. In the other infection systems that we characterised through this approach (hepatocyte/hepatitis C virus ^57^ and mosquito cells/*Wolbachia* endosymbiont ^58^) the functional relevance of the most interesting array hits was established through siRNA campaigns. In the erythrocyte/*Plasmodium* system where siRNA-based knock-down of host genes is not feasible, we propose that similar Western blot validation should be undertaken as the first step in further work on any of the numerous additional enzyme/pathways identified as hits in the array.

The need for systematic validation notwithstanding, the extent to which host erythrocyte signaling is modulated by *Plasmodium* infection is surprising. The array data are consistent with earlier observations implicating some of the host cell signaling pathways in the infection process, and allowing an in-depth, highly specific understanding of the mobilisation of such pathways (for example, see above the section pertaining to PKC isoforms). The observation (Fig. 4c) that B-Raf is present at the ring-stage but appears to be degraded in trophozoite- and schizont-infected RBCs is consistent with a thorough modulation of host erythrocyte signaling pathways during infection, not only through activation/inactivation of enzymes by phosphorylation, but also through affecting the abundance of specific signaling elements. In line with this observation, the host erythrocyte cAMP-dependent kinase PKA has been reported to similarly be degraded in infected cells^59^. This is reminiscent of the strategy of other intracellular pathogen to modify the composition of signaling pathways in their host cell. A well-characterised example is that of the Lethal Factor, a component of the anthrax toxin that is secreted by *Bacillus anthracis* and that possesses highly selective endopeptidase activity directed against host cell MEKs, thereby inactivating MAPK pathways (reviewed in ^60^). Down-regulation of MAPK pathways through degradation of specific components is not restricted to host-pathogen interactions. For example, during hematopoietic cell apoptosis, IL-3-deprivation results in the cleavage of Raf-1 by caspase 9, causing separation of the N-terminal regulatory domain from the C-terminal kinase domain^61^. Likewise, c-MET and several receptor tyrosine kinases^62^ can undergo specific proteolytic cleavage events, which mediate various biological outcomes (reviewed in ^63^). It is therefore interesting to note that in our Western blot analysis, c-MET appears as a band of lower mass in trophozoite/schizont RBCs than at the ring stage (Fig. 5). It is tempting to speculate that intracellular pathogens hijack such cellular processes to suppress host pathways. Our observation that host erythrocyte B-Raf is activated early in infection (ring-stage), and apparently subsequently degraded, warrants further studies that will yield fascinating insights into host-parasite interactions. In particular, it will be of great interest to investigate if some host pathways identified as activated in the present study are dependent on the replacement of specific host cell kinase by parasite-encoded kinases, several of which are known to be exported to the host erythrocyte (e.g. ^64,65^).

A major long-term objective of this research is the identification of antimalarial compounds with untapped modes of action. It is a clear possibility that part of the erythrocyte signaling response to infection may be directed towards activating innate immune defence mechanisms, and thus downregulating such responses would be undesirable. However, based on the hypothesis that at least some of the signaling events triggered in the host cell by infection are required for parasite survival, we tested highly selective inhibitors against some of the activated kinases and showed that these compounds have high potency (in the low nM range) in *in vitro* culture systems. The nanomolar activity against both *P. falciparum* and the phylogenetically distant *P. knowlesi* for c-Met and B-Raf inhibitors indicate that the reliance on the activation of host erythrocyte pathways spans various *Plasmodium* species. Importantly, the markedly impaired proliferation of *P. berghei* in a murine malaria model treated with the c-Met inhibitor PHA-665752 shows that targeting host kinases to control malaria infection *in vivo* is achievable, strengthening the case for kinase-focussed host-targeted intervention.

Major attractive features of targeting host-cell kinases as a strategy to develop novel antimalarials are threefold: First is the effectiveness of specific inhibitors across phylogenetically distant *Plasmodium* species, illustrated by the facts (i) that inhibitors against three host cell kinases activated by infection (MEK1, c-Met and B-Raf) display similar *in vitro* potency on *P. falciparum* and *P. knowlesi*, and (ii) that MEK and c-Met inhibitors also have activity against the *P. berghei* rodent malaria parasite *ex vivo* (MEK ^15^) and *in vivo* (c-MET). A second tremendous advantage of this strategy is that the most parsimonious pathway to resistance, i.e. the selection under drug pressure of genotypes encoding a mutated target with decreased susceptibility, is not possible if the target is encoded by the host. Third, large kinase-directed compound libraries are available from the intensive efforts made over the past two decades in the context of cancer chemotherapy (including inhibitors of many of the kinases identified in this study as being activated in infected erythrocytes), providing a unique repository of potential antimalarials. These can now be exploited as a basis for fundamental research into host-parasite interactions, and as leads for the development of novel antimalarials with untapped modes of actions.

## Methods

### *Plasmodium* spp. culture and life stage synchronisation

Human erythrocytes were supplied by the Australian Red Cross. *P. falciparum* (clone 3D7) and *P. knowlesi* (YH1) were grown in human erythrocytes as previously described ^66–69^. *P. falciparum* synchronisation (4-hr window) was achieved using 5% w/v Sorbitol as well as 15 μl/ml culture of heparin ^70^. Both techniques were utilised for the trophozoite and schizont stage samples used on the antibody microarray, whereas an 8-hr synchronisation window was achieved through sorbitol alone for the ring stage. Purification of *P. falciparum* trophozoite and schizont-infected cells was achieved using a super-MACS column ^71^. *P. knowlesi* cultures for *in vitro* assays were partially synchronised (~12-hr age range) using heparin ^67^.

### Kinexus antibody microarray preparation

The Kinex 900P array kits were purchased from Kinexus. Protein extracts were prepared as per the manufacturer’s instruction. Each array was, unless otherwise specified, loaded with protein extracts normalized to equivalent cell number using a hemocytometer. Array scanning was conducted by the manufacturer at their facility. Arrays were loaded with protein extracts of rings 4-12 hrs (approximately 33% parasitemia, n=3), trophozoites 24-28 hrs (magnet purified, n=3) and schizonts 44-48 hrs (magnet purified, n=2). Parallel uRBC control extracts containing RBCs from the same donor as the iRBC treatment were loaded onto the arrays to measure baseline RBC phosphorylation (these cells were cultured in the same conditions as the parasite-infected samples for approximately 24 hours before harvesting). To identify phosphosites of parasite proteins cross-reactive to the array, saponin pellets and supernatants of magnet purified, saponin lysed trophozoites/schizonts were compared. Saponin pellet samples were loaded at 2 mg/ml on the array, 20x more pellet material than is present in 2 mg/ml of equivalent saponin supernatant material.

### Western blotting

Western blot analysis of infected and uninfected erythrocytic material performed on protein extracts prepared by resuspending cells in M-PER mammalian protein extraction reagent (Pierce) supplemented with 1x protease inhibitory cocktail (EDTA free) (Roche), 300 mM benzamidine, 200 mM phenylmethylsulfonyl fluoride, 500 mM sodium fluoride, 100 mM sodium orthovanadate and 500 mM β-glycerophosphate. Lysates were cleared by centrifugation (10,000 g for 15 mins at 4°C); samples were heated (70°C) in Laemmli reducing sample buffer before separation on 4-12% Tris/Bis gradient SDS-PAGE gels (Invitrogen). After electrophoresis, proteins were transferred to nitrocellulose membrane (Amersham) and blocked (1-hr at room temperature) with 5% Skim milk (Diploma) in Tris-buffered saline containing 0.05% Tween 20 (TBST). Immunoblotting was performed using the following antibodies; phosphorylated c-MET (Y1234/Y1235; D26, Cell Signaling Technologies), pan c-MET (D1C2, Cell Signaling Technologies), pan B-Raf (OTI4B2, Bio-Rad), phosphorylated B-Raf (S729, AB-PK535, Kinexus), Glycophorin-C (Abcam). All primary antibodies used at 1:500 dilution and incubated overnight at 4°C. Anti-Rabbit/Mouse-horseradish peroxidase-conjugated secondary antibody was used for all experiments at 1:2500 dilution (Monoclonal antibody facility Monash University Clayton, Vic, Australia).

### IC_50_ determination

*P. falciparum* – The IC_50_ values of Crizotinib, PHA-665752, SB590885, Sorafenib and Artemisinin were determined with asynchronous cultures with a starting parasitemia of 0.25% in 2% haematocrit. Cultures were incubated with the compounds for 72 hrs before adding SYBR-gold nucleic acid stain (1:10000 dilution) (ThermoFisher Scientific) modified from ^72^ substituting SYBR-green for SYBR-gold. Plates were incubated with SYBR-gold for 1 hr in the dark prior to reading fluorescence on a Tecan plate reader (I-control software), and the inhibitory IC_50_ determined using GraphPad PRISM (GraphPad software).

*P. knowlesi* – *P. knowlesi* growth inhibition assays using ring-stage parasites were set up at 1% parasitaemia and 1% haematocrit in 96-well round-bottom plates at a final volume of 45 μL as described ^73,74^. A 10x final concentration of drug and controls was added to make the final volume to 50 μL and the drug assay cultured for 50 hrs until parasites reached late trophozoites in the next growth cycle. Assays were stained with 10 μg/mL ethidium bromide (EtBr, Bio-Rad) for 1hr and washed prior to flow cytometry (Becton Dickinson LSR) assessment of parasitaemia with gating as per established protocols ^75^. Parasitaemia counts were quantitated using FlowJo software (Tree Star) and the inhibitory IC_50_ determined using GraphPad PRISM (GraphPad Software).

### Time-dependent inhibitor treatment of parasites monitored via flow cytometry

Highly synchronous cultures (0 – 4 hrs post-invasion) at 2% haematocrit treated with DMSO (vehicle), Crizotinib, PHA-665752, SB-590885 or artemisinin at 5x the IC_50_ values were monitored at 12-hr time intervals over two full asexual intra-erythrocytic cycles (84 hrs) by flow cytometry. Live parasitaemia was quantitated by a dual-colour flow cytometry staining protocol using 2 μM Hoechst-33342 staining for 8 minutes and 75 nM MitoTracker Orange for 25 minutes. Staining was completed in v-bottom plates with two washes in complete RPMI. Stained cells were transferred to polypropylene tubes and diluted one in four in complete RPMI. Cells were then immediately analysed on an LSR BDFortessa™ with the laser UV379 and filter 450/50 for Hoechst-33242 staining, and laser YG585 with filter 585/15 for MitoTracker Orange staining. Biological replicates (n=5) were normalised to a 2% total parasitemia value at the first-time point (0 - 4 hrs post-invasion).

### *P. falciparum* stage-dependent inhibitor treatment and washout out assay

*P. falciparum* cultures were synchronised into a 6-hour development window at 2% haematocrit and treated with 5x the 72hour IC50 of SB590885, Artemisinin or DMSO at 0-6hpi (ring stage treatment) 22-28hpi (trophozoite stage treatment) and 38-44hpi (schizont stage treatment). Treatments were washed from the respective cultures after 4 hours (denoted by the black arrow) and monitored until the next cycle of parasite development to assess parasite survival using flow cytometry (2 μM Hoechst-33342 staining for 8 minutes). Starting parasitemia was normalised to 5% across the biological replicates (n=3) error bars represent the standard error of the mean.

### *P. berghei in vivo* drug treatment assay

Animal research was conducted under licenses from the UK Home Office, and protocols were approved by the Animal Welfare and Ethical Review Body of the Wellcome Sanger Institute. Mice were purchased from Envigo, kept in specific-pathogen-free conditions and subjected to regular pathogen monitoring by sentinel screening. They were housed in individually ventilated cages furnished with autoclaved aspen woodchip, fun tunnel and Nestlets at 21°C +/- 2°C under a 12:12 hr light-dark cycle at a relative humidity of 55 +/- 10%. They were fed a commercially prepared autoclaved dry rodent diet and water, both available *ad libitum*. The health of animals was monitored by routine daily visual health checks. Cohorts of seven female, 10-week-old BALB/c mice per treatment group were infected by i.v. inoculation with 10^6^ infected erythrocytes. Assays to determine parasite growth were performed as described ^76^, using the reporter parasite line PbGFPLuccon (RMgm-29 in the rodent malaria genetic modification database, http://www.pberghei.eu/index.php?rmgm=29), which expresses a GFP-firefly luciferase fusion protein under the control of the constitutive *eef1a* promoter ^77^. Treatment with PHA-665752 or solvent control was 3hr thereafter and on day two post infection by i.p. injection of 25 mg/kg/day of PHA-665752. The drug was prepared fresh by diluting a stock of 50 mg PHA-665752 in 0.5 ml DMSO (w/v) in water (1:40, v/v). The same drug formulation and treatment regime had proved effective in controlling carcinoma growth ^56^.

## Data Availability Statement

All raw data pertaining to this study are available in Supplemental Datasets 1 to 3.

## Acknowledgements

We thank Dr Kylie Quinn for lending her Flow Cytometry expertise and the staff of Monash FlowCore Facility, in particular Andrew Fryga and Adam Dinsdale, for their continued experimental assistance. We also thank Prof Roger Daly, Dr. Teresa Carvalho, A/Prof Jose Garcia-Bustos and Prof Brian Cooke for fruitful discussions. This work was supported by the Australian Government National Health and Medical Research Council (Project Grants APP1082619 [CD] and APP1143974 [DW]), Australian Research Council PhD Scholarship (AB), Hospital Research Foundation Fellowship (DW) and internal support from Monash University and the University of Adelaide (Beacon Fellowship (DW).

## Notes

### Competing Interest Statement

The authors have declared no competing interest.

