## Supplementary figures and images for "Signalome-wide assessment of erythrocyte response to *Plasmodium* reveals novel targets for host-directed antimalarial intervention"

### Supplementary Figure 1

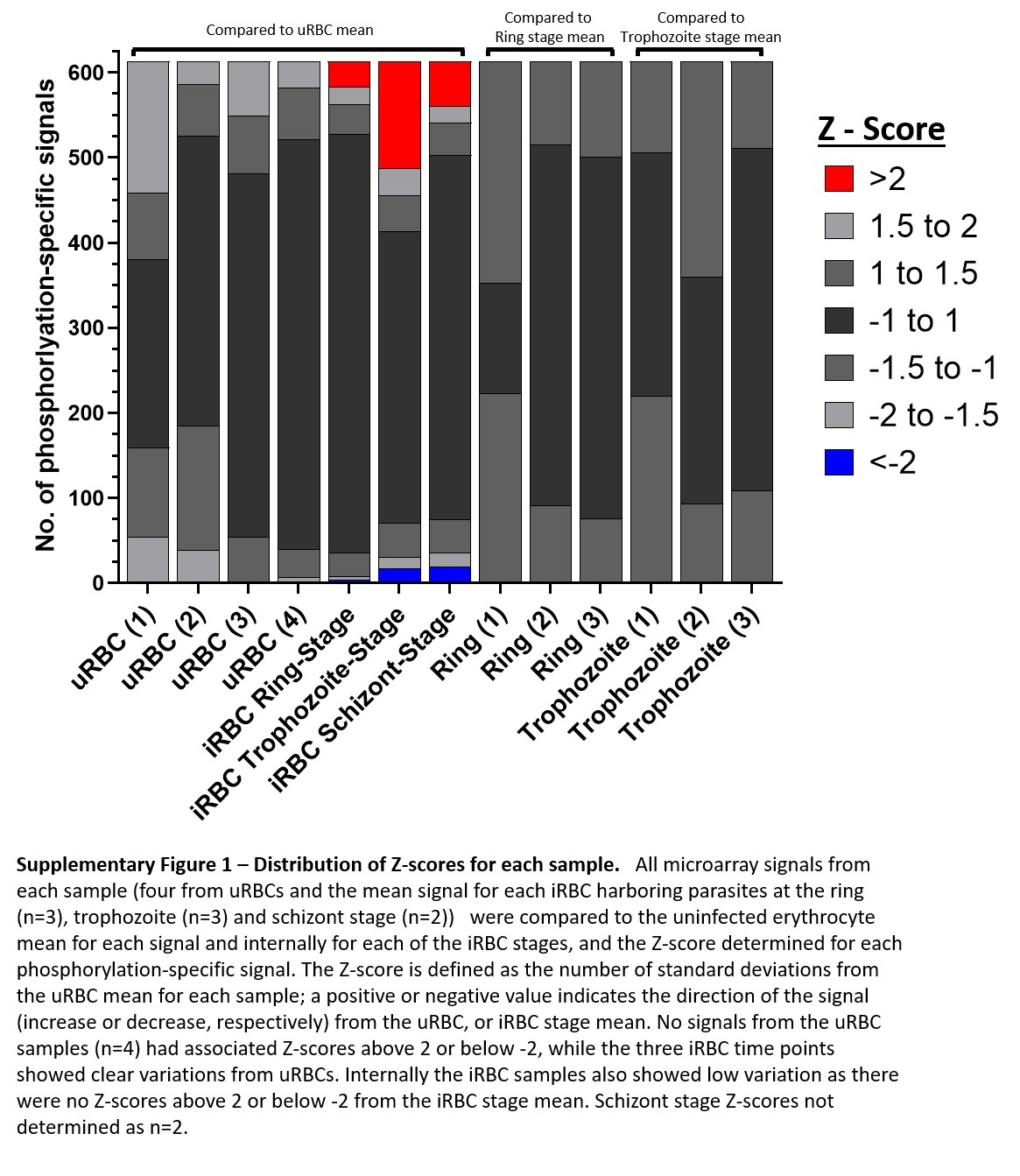

### Supplementary Figure 2

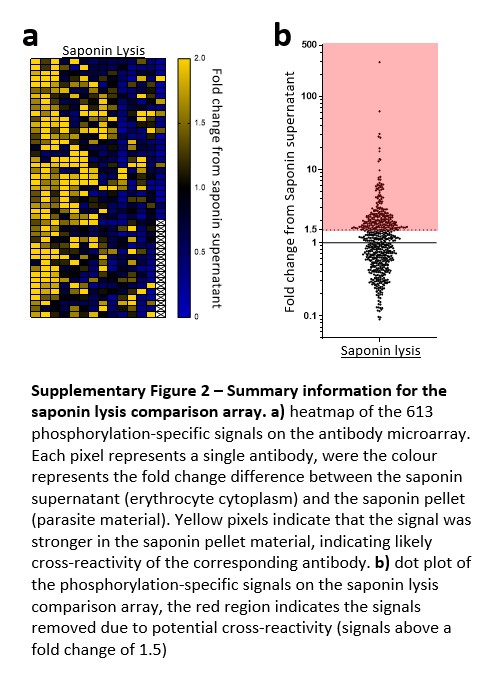

### Supplementary Figure 3

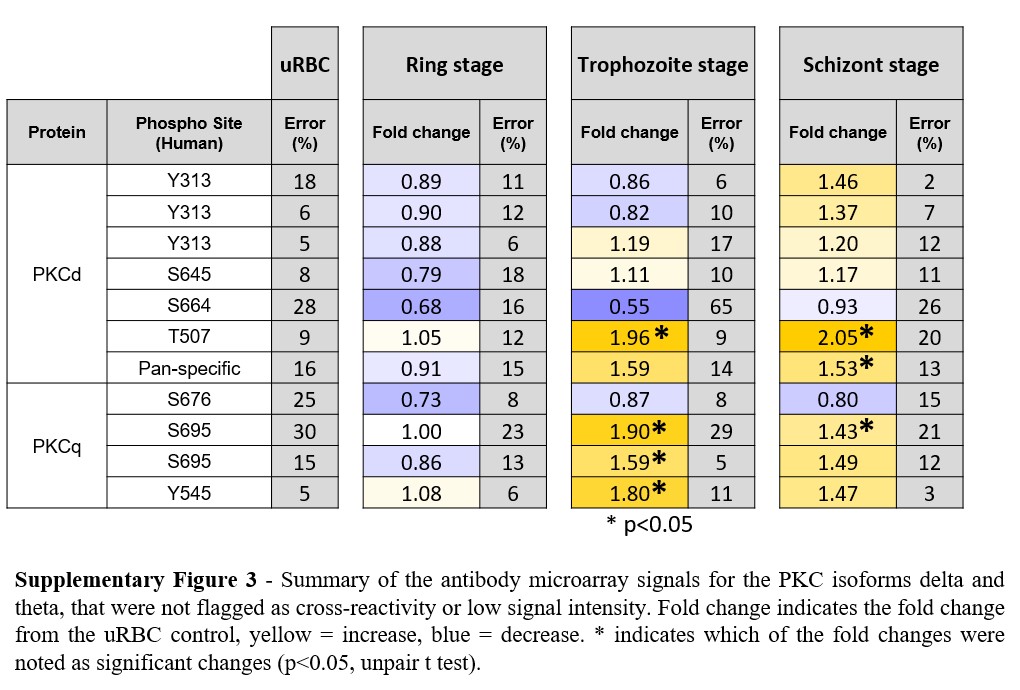

### Supplementary Figure 4

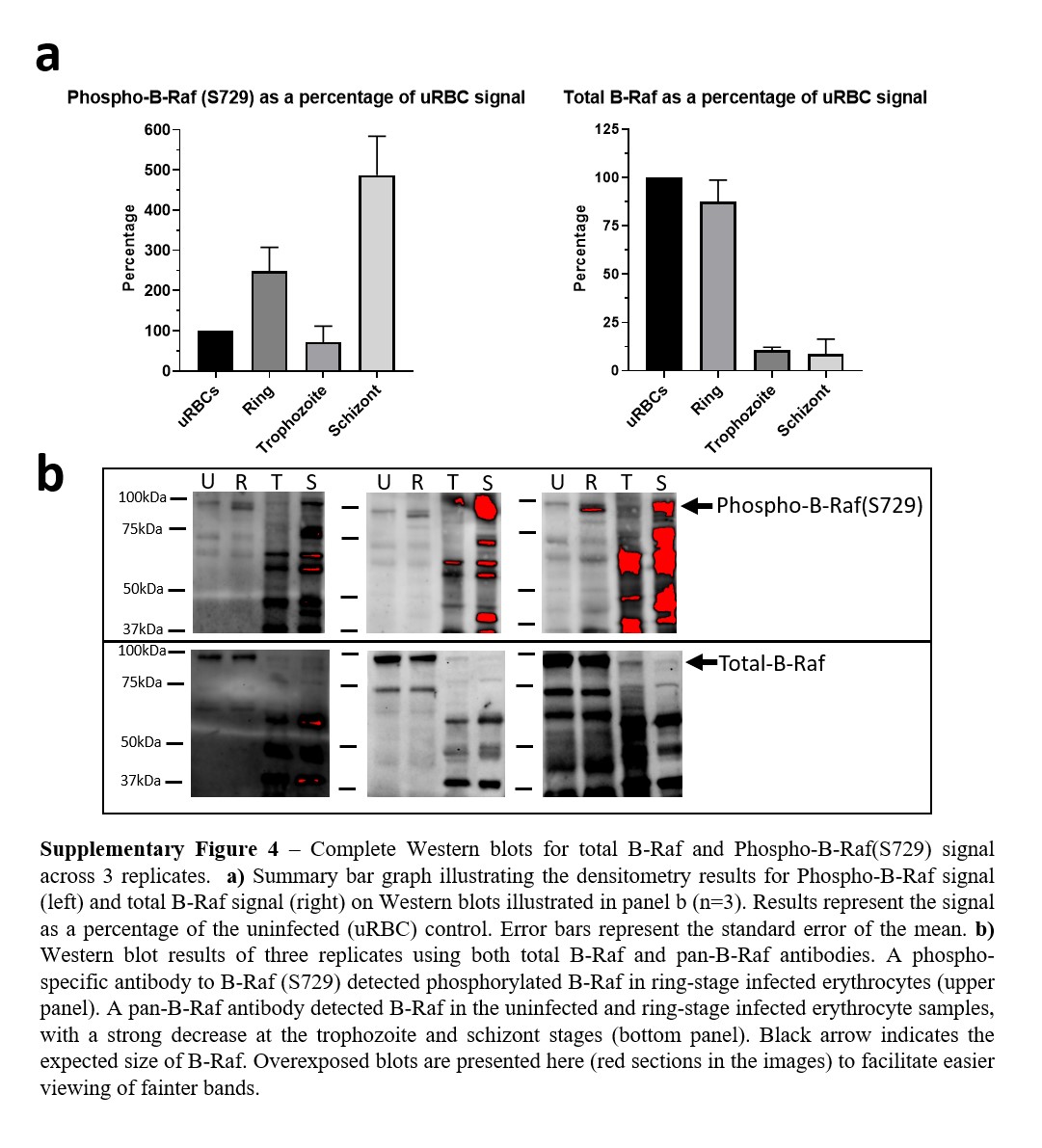

### Supplementary Figure 5

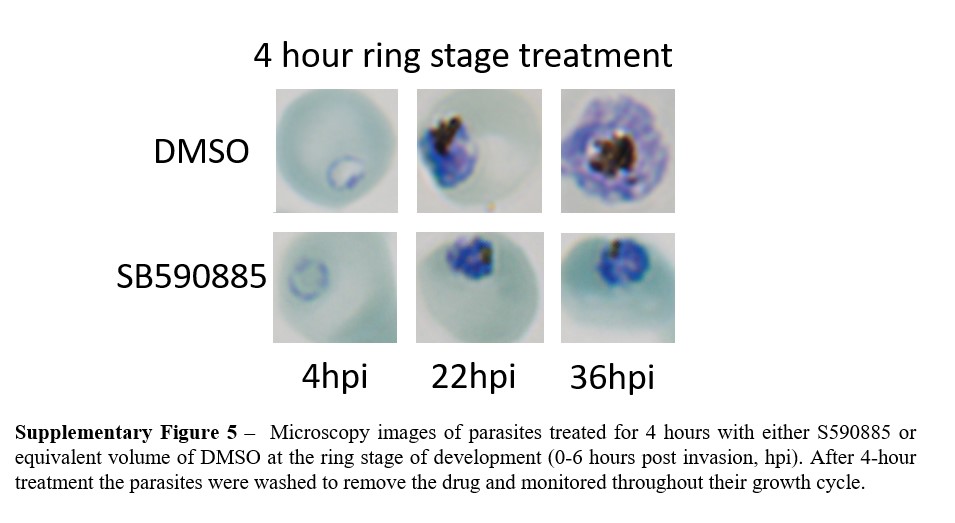
